# modleR: a modular workflow to perform ecological niche modeling in R

**DOI:** 10.1101/2020.04.01.021105

**Authors:** Andrea Sánchez-Tapia, Sara Ribeiro Mortara, Diogo Souza Bezerra Rocha, Felipe Sodré Mendes Barros, Guilherme Gall, Marinez Ferreira de Siqueira

## Abstract

Ecological niche models (ENM) use the environmental variables associated with the currently known distribution of a species to model its ecological niche and project it into the geographic space. Widely used and misused, ENM has become a common tool for ecologists and decision-makers.

Many ENM platforms have been developed over the years, first as standalone programs, later as packages within script-based programming languages and environments. The democratization of these programming tools and the advent of Open Science brought a growing concern regarding the reproducibility, transparency, robustness, portability, and interoperability in ENM workflows.

ENM workflows have some core components that are replicated between projects. However, they have a large internal variation due to the variety of research questions and applications. Any ecological niche modeling platform should take into account this trade-off between stability and reproducibility on one hand, and flexibility and decision-making on the other.

Here, we present **modleR**, a four-step workflow that wraps some of the common phases executed during an ecological niche model procedure. We have divided the process into (1) data setup, (2) model fitting and projection, (3) partition joining and (4) ensemble modeling (consensus between algorithms).

**modleR** is highly adaptable and replicable depending on the user’s needs and is open to deeper internal parametrization. It can be used as a testing platform due to its consistent folder structure and its capacity to control some sources of variation while changing others. It can be run in interactive local sessions and in high-performance or high-throughput computational (HPC/HTC) platforms and parallelized by species or algorithms. It can also communicate with other tools in the field, allowing the user to enter and exit the workflow at any phase, and execute complementary routines outside the package. Finally, it records metadata and session information at each step, ensuring reproducibility beyond the use of script-based applications.

## 1 Introduction

Ecological niche models (ENM) use the currently known distribution of a species and the environmental variables associated with it to model their ecological niche in the environmental space and project it into the geographic space. ENM have been used to bring up and test biogeographic hypotheses, or in applications regarding conservation biology, invasion biology, among others, and has become a basic tool for ecologists and decision makers.

ENM workflows have a general structure that is relatively common to all projects, and internal variation that depends directly on the research question and guide decision-making for the variety of possible applications. The specific parametrization and level of detail of each step depend and should reflect the research questions and hypotheses, and there is not a common recipe that will suit every project. However, a minimal ENM workflow is expected to include some basic steps. These steps include:

- The inspection of occurrence data quality, data cleaning, the selection of biologically meaningful, uncorrelated environmental variables, with a resolution and extent matching the quality of the occurrence data (Giannini et al., 2012).
- The delimitation of the calibration area – that is supposed to reflect the accessible areas for the species, i.e. areas that have been explored by the species and found appropriate or not (Peterson et al., 2011; Barve et al., 2011).
- Data partitioning into test and training sets to perform model fitting and validation (Fielding and Bell, 1997), i.e. checking the model’s predictive ability.

Some other steps are frequent but optional. For example, following the modeling procedure, and depending on the application, the algorithms with the best performance may be chosen or combined in an ensemble model *sensu* Araújo and New (2007) (i.e. algorithmic consensus models).

Since their conception, a large number of platforms to perform ENM have been created, such as Maxent *(Phillips et al., 2006), DesktopGarp (Scachetti-Pereira, 2002)*, Ecological Niche Factor Analysis (ENFA, Hirzel et al. (2002), diva-gis ^1^, and openmodeller (de Souza Muñoz et al., 2009). These were initially mostly point-and-click standalone applications and focused on model fitting and projection. Other phases of the whole ENM process, such as data preparation or output processing, were performed in a variety of tools.

With the advent of Open Science as a philosophy to guarantee the access, robustness, transparency and reproducibility of the scientific workflows, the use of script-based programming languages and environments in ENM is being favored. The R statistical environment (R Core Team, 2018) is one of the current script-based environments that allow for such reproducible applications, due to its large collection of functions and packages. Today, even Maxent, the most widely used ENM application, has been implemented in R via package **maxnet** (Phillips, 2017), which facilitates its integration into the whole R ecosystem. Among the common current reproducible ENM applications in R, we may cite now classic packages such as **BIOMOD2** (Thuiller, 2003), **dismo** (Hijmans et al., 2017), **and ENMTools** (Warren et al., 2019), and equally important newer frameworks such as **sdm** (Naimi and Araújo, 2016), **ENMeval** (Muscarella et al., 2014), **Wallace** (Kass et al., 2018), **and zoon** (Golding et al., 2017). In addition to these, many groups are developing tools to assist in one or more parts of the workflow (**spThin** Aiello-Lammens et al. (2015), **kuenm** Cobos et al. (2019), **blockCV** Valavi et al. (2019)), **among others**.

Today, R must be understood as the platform that can integrate different ENM applications. For this, developers should be concerned with integration among packages, which implies in having modular structures. Workflows need to be able to expand as new techniques are tested and validated, and consensus regarding each step is reached. We also need ENM workflows that take into account the trade-off between workflow stability (that all phases are duly executed) and decision-making (variety of applications and parametrization).

In addition to this, it is important to note that the use of script-based applications does not guarantee reproducibility *per se*. Data processing, decision steps and parametrization options should be documented as metadata. Feng et al. (2019) listed 33 steps to check the reproducibility of ENM workflows, grouped in (A) occurrence data collection and processing, (B) environmental data collection and processing, (C) model calibration and (D) model transfer and evaluation By using script-based analysis combined with detailed metadata, workflows in ecology can achieve reproducibility and transparency (Powers and Hampton, 2019). Thourough documentation also means saving the subproducts from each step, so that they can be examined, corrected, reused, and reinterpreted.

Here, we present **modleR**, a workflow designed to automatize some of the common steps when performing ecological niche models, using the R statistical environment (R Core Team, 2018). An early version (Sánchez-Tapia et al., 2018) had a similar structure but several improvements have been implemented to this day, so we refer the users to the current manuscript, with focus on the R package. We have used functions from well-known packages, such as **dismo** (Hijmans et al., 2017), **maxnet** (Phillips et al., 2017), **RandomForest** (Liaw and Wiener, 2002), and some newer but promising implementations, such as **kuenm** (Cobos et al., 2019). The workflow is highly adaptable and replicable depending on the user’s needs, and is open to deeper internal parametrization. In order to communicate with other ENM tools within the R environment, it does not create new object classes or methods In addition to this, **modleR** records metadata and session information (i.e. loaded packages, their version and origin) for each step in the modeling procedure in a structured way. Our workflow complies with all the procedures listed by Feng et al. (2019) regarding model calibration and model transfer and evaluation (Table 1).

**Table 1:** Reproducibility information provided by **modleR** following the checklist by Feng et al. (2019).

| Checklist | Category | Item | Where to find |
| --- | --- | --- | --- |
| **Model calibration** | Data input | Extent of background data | data_setup/metadata.csv |
|  |  | Number of background data | data_setup/metadata.csv |
|  |  | Sampling method for background data | data_setup/metadata.csv |
|  |  | Variable selection | data_setup/metadata.csv |
|  | Algorithm | Algorithm name | partitions/metadata.csv |
|  |  | Version of algorithm and software | partitions/sessionInfo.txt |
|  |  | Parameterization | partitions/metadata.csv |
| **Model transfer and evaluation** | Evaluation | Evaluation index | final_model/metadata.csv |
|  |  | Threshold for evaluation index | final_model/metadata.csv |
|  |  | Dataset used to evaluate models | data_setup/metadata.csv |
|  | Output | Format/transformation | partitions/metadata.csv |
|  |  | Threshold | partitions/metadata.csv |

**modleR** keeps memory usage low by saving outputs to the hard disk and uses a nested folder structured that can be used as a platform for performing experiments or tests. Finally, it can run locally, or on high performance or high throughput computational (HPC/HTC) platforms, and be parallelized easily within the R environment. Throughout the text, we explain the theoretical basis for some of the parametrization and highlight best practices to ensure the robustness and reproducibility of these analytical workflows.

## 2 Workflow structure

### 2.1 Workflow overview

**modleR** workflow consists of mainly four steps that could be used sequentially (Figure 1). Each step is self-contained, allowing the user to enter or exit the workflow at any step.

**Figure 1:**
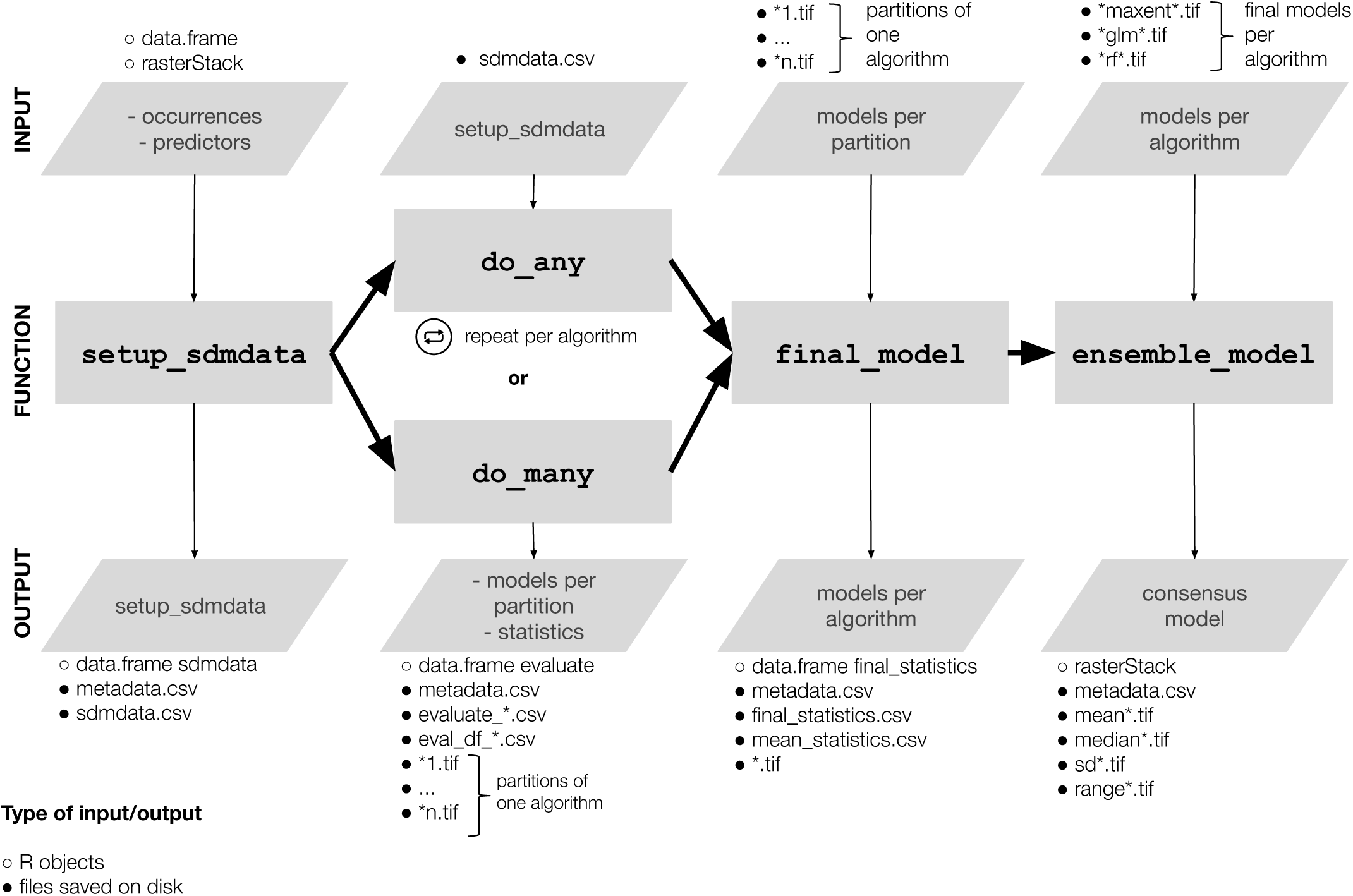
Overview of the **modleR** workflow steps

#### Data setup

Given the occurrence records and a set of environmental predictors, the first step of the workflow prepares the biotic data by applying some data cleaning and geographic filters, and assists in the selection of variables by controlling their correlation. It incorporates several options for defining the calibration area and including real absence data or sampling pseudoabsences. Furthermore, it executes the data partition using k-fold cross-validation or bootstrap procedures.

#### Model fitting and projection

The second part of the workflow fits the ecological niche models for each of the partitions defined in the first step, and projects them into the geographical space, using algorithms implemented in the R statistical environment. This step also evaluates the models, and returns summaries of the performance metrics along all the threshold values and for given specific thresholds. Optionally, it also projects the models into other environmental datasets (for the past or the future, or different geographical locations, for example).

#### Partition joining

The third part of the workflow presents several options to join the partitions fit for each algorithm into one final model per species per algorithm and to visualize the variation between partitions.

#### Ensemble modeling - algorithmic consensus

The fourth part of the workflow allows the user to select the best performing algorithm or to create an algorithmic consensus model using different methods, and to visualize the variation between algorithms.

By setting these steps in an sequential way, **modleR** constitutes an ecological niche modeling platform that allows for repeated modeling rounds in a controlled environment. Because the modular workflow is build to enhance reproducibility, at all steps a metadata file is written with all main parameter values and associated information of the ENM workflow.

### 2.2 Output definitions and folder structure created by modleR

#### 2.2.1 Output definitions

Throughout the text, we define the output of each step of the workflow as follows (Figure 2.2.1):

**Figure 2:**
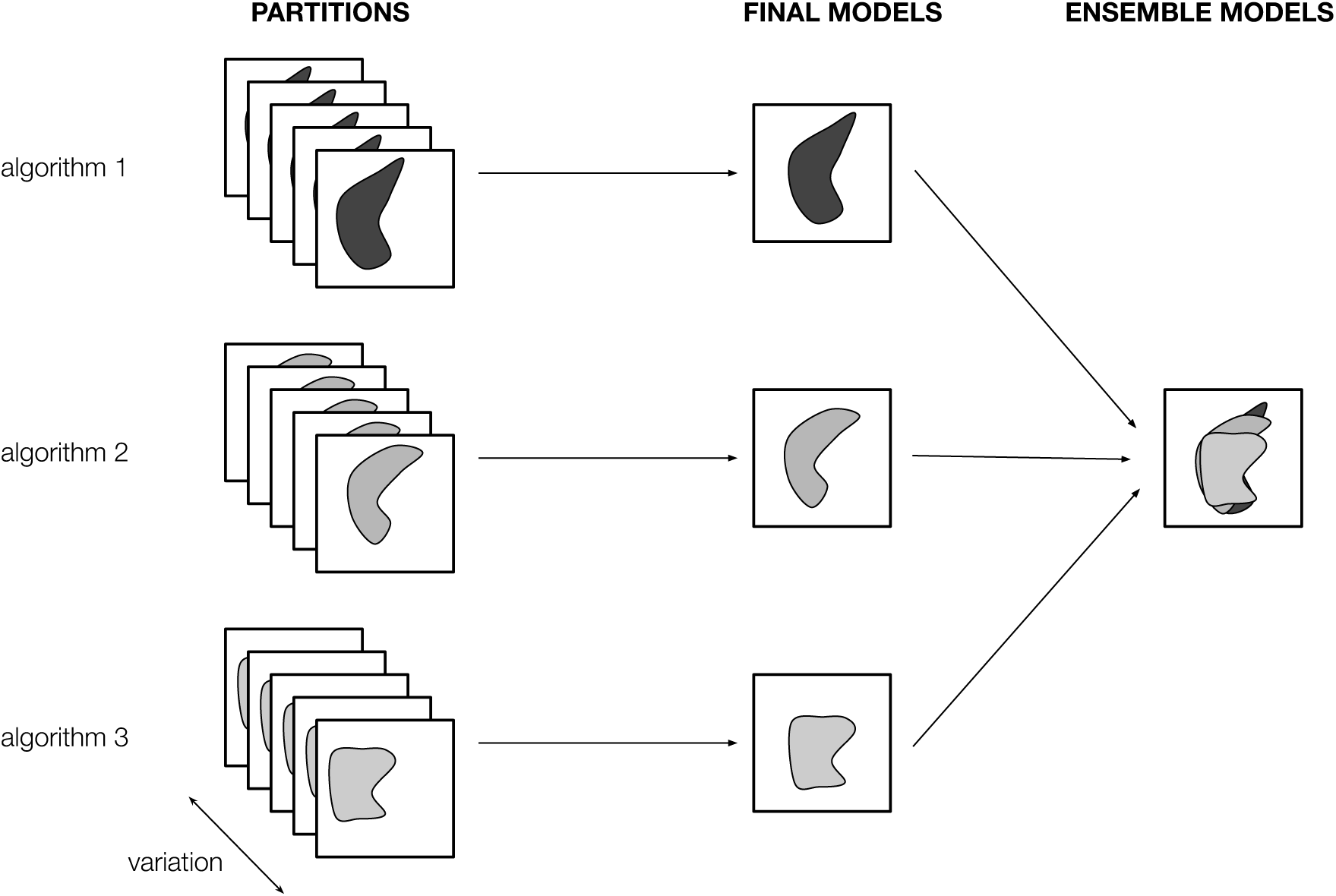
Partition, final models, and ensemble models in the **modleR** workflow

- A **partition** is the result of the individual modeling round, that takes one training and test dataset and a single algorithm. Although a single partition can be created, the usual is performing several modeling rounds. A bootstrap will create several partitions through resampling, a k-fold crossvalidation will create k partitions, and a repeated (n times) k-fold crossvalidation will create n * k partitions. Partitions refer to present conditions and to models projected to other environmental variables datasets.
- A **final model** results from joining together the partitions and obtaining *one model per species per algorithm*.
- An **ensemble model** is the process of joining together the results obtained by different algorithms, to obtain *a single algorithmic consensus model per species* (Araújo and New, 2007).

#### 2.2.2 Folder structure

**modleR** was built to write on disk the outputs of each step as described below. This is done for three main reasons. First, it keeps memory usage low. Second, it allows the user to enter and leave the workflow at any step, to continue in a different session at any time or use other tools in the R environment. Finally, an easily readable folder structure both to the user and the computer is essential for reproducibility and portability.

- A whole modeling round will be created in a single folder - by default `models_dir = “./models_dir”`, that is, a folder within the current working directory, as indicated by the period.
- Each species will be saved in its own subfolder, `“./models_dir/Genus_epithet”` by default. **modleR** will format species names in order to avoid non-ascii characters and spaces.
- Inside each species subfolder, a folder for each projection will be created. The default (and non-optional) projection to the present conditions is called `present`, but additional folders will be created for each projection and named after each one (**see section 4**.**3**).
- Within each projec
- Within each projection folder, the following folders will be created, corresponding to the four steps of the workflow

– **data setup** (only for the projection in the present)
– **partitions**
– **final models**
– **ensemble**

The resulting folder structure is shown in Figure 3.

**Figure 3:**
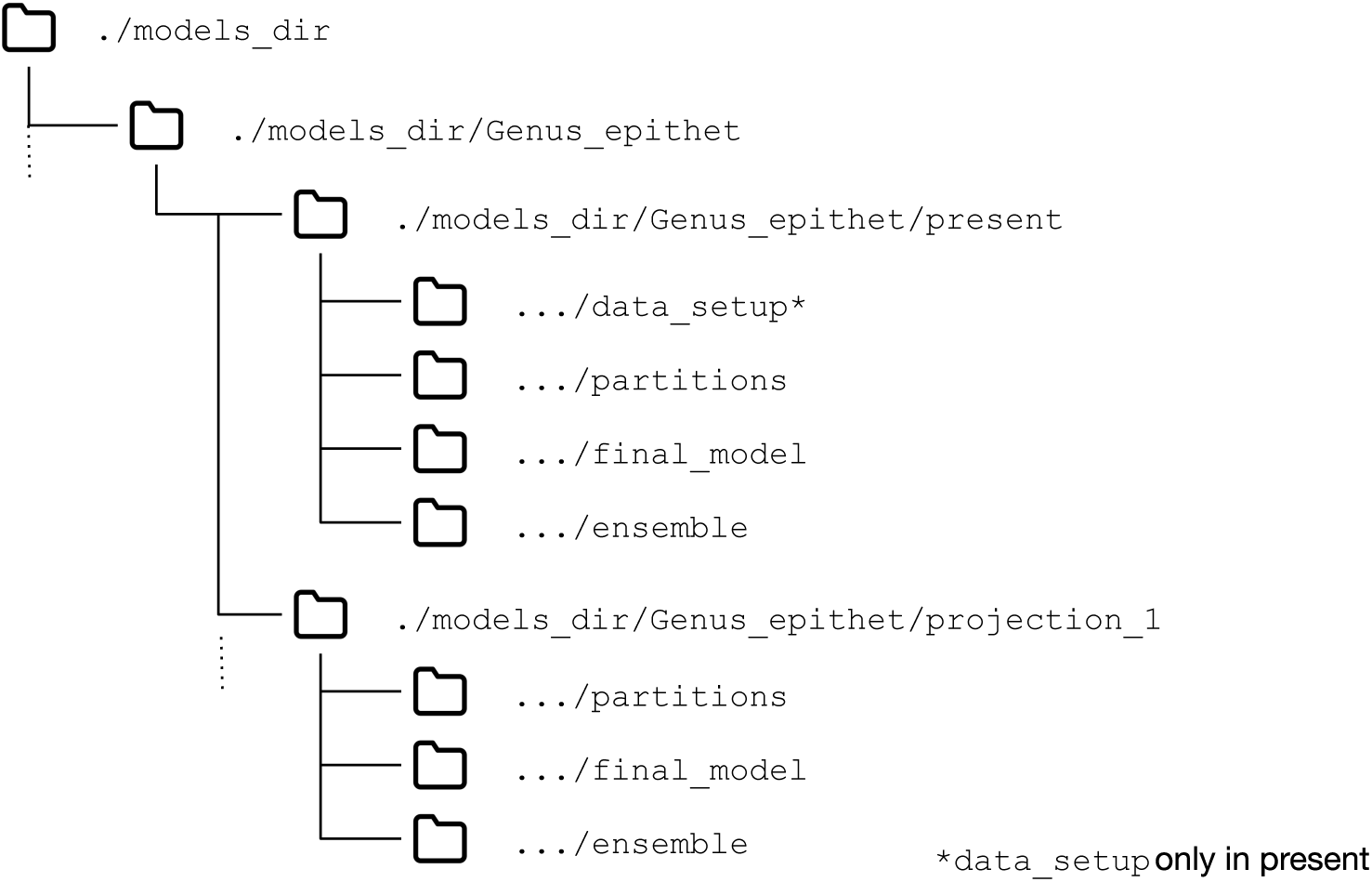
folder structure created by **modleR**

Parameter `models_dir` allows to set the output folder anywhere in the hard disk. For reproducibility purposes, it is recommended that this folder be referred as a relative path as in the default value (e.g. `“./models_dir”`) rather than as an absolute folder (e.g. `“C://Documents/…/models_dir”`), that will not be portable to different computers. The *names* of the final and ensemble folders can also be modified, but the nested subfolder structure will remain the same. If the user changes any of these names, they will need to include the new value when calling for the rest of the functions. We strongly recommend not using non-ascii characters or spaces on folder names.

This partial flexibility allows for experimenting with final model and ensemble construction. For example, final or ensemble can be run more than once in different output subfolders, with different parameters or subsets of the algorithms that were fit initially.

In the following sections we will detail the options and implementation decisions of each of the workflow steps.

## 3 Data setup: setup_sdmdata()

Function `setup_sdmdata()` performs basic data cleaning, data partitioning into training and test sets, and pseudoabsence sampling for model calibration to take into account the geographic area accessible to the species. It also assists in variable selection according to an upper correlation limit within the calibration area. This step writes on disk the data needed to fit ENM models in further steps of the workflow, and the associated metadata and session information, including the version and origin of the packages used.

### 3.1 Cleaning and thinning occurrence data

Function `setup_sdmdata()` will eliminate exact duplicates, occurrences with no environmental data and occurrences in the same pixel of the environmental predictors through arguments `clean_dupl`, `clean_nas` and `clean_uni`, respectively. We assume that taxonomic and geographic cleaning were performed by the user. Data cleaning inside our package is mostly directed to fit observations to the predictor variables’ scale and algorithm requirements.

For instance, some algorithms can receive only presence data and will later run even with NAs whereas some others cannot run with NAs in the data table. By default all data cleaning procedures are set to `FALSE` to avoid computational costs and assuming that the user is responsible for this step. The user may clean the data outside the application, either within R or using dedicated software that records the cleaning steps, such as OpenRefine ^2^.

In addition to this, the function can thin the occurrences according to a spatial grid, using parameter `geo_filt`. This step intends to control the sampling bias in occurrence data (Varela et al., 2014) and could also be performed by controlling the environmental similarity of the dataset. In **modleR**, this step is implemented according to Varela et al. (2014), within a grid, but there are other methods, packages and functions in R that can perform geographic and environmental thinning, of which we highlight **spThin** (Aiello-Lammens et al., 2015).

### 3.2 Pseudoabsence selection options

The next step regarding data setup is the definition of absences or pseudoabsences within calibration areas in order to perform model fitting and data evaluation. As a first option, **modleR** can receive user-defined real absences (`real_absences = TRUE`) as a dataframe object with longitude and latitude columns. In this case, no pseudoabsence will be sampled internally.

In the most likely case that real absences are not available, pseudoabsences must be sampled in order to calibrate and evaluate the models. When sampling for pseudoabsences, the number, sampling scheme and minimum and maximum distance from occurrence points affect considerably model performance and evaluation (Barbet-Massin et al., 2012), and **modleR** has several options in this regard.

The number of pseudoabsences can be set using parameter `n_back`. However, when applying maximum and minimum distance buffers for sampling (see next), the final sampling area may not be large enough to fit the number of pseudoabsences requested by the user, so the function will check internally that enough unique pixels are available in the calibration area and reset to a maximal n_back value. If this modification takes place, the metadata (in `metadata.csv`) will reflect it.

The spatial configuration for pseudoabsence sampling is another key element for pseudoabsence sampling, because it should reflect the accesible area for the species, i.e. areas that were most likely explored by the species and found unsuitable (Barve et al., 2011; Peterson et al., 2011; VanDerWal et al., 2009), so it should not be performed in areas too far away from the occurrences.

We have implemented a scheme for pseudoabsence sampling, with two steps that can be used alone or simultaneously (Figure 4). The first step is an inclusion buffer that sets the maximal geographical boundaries for pseudoabsence sampling, so only accessible areas for the species are taken into account, and the second step is an exclusion filter, that sets the minimum distance for pseudoabsence sampling, so that areas too close (either in the geographical or the environmental space) to the occurrence points are omitted from sampling. These steps have been used to improve the model’s discrimination ability and to control for spurious good performance.

**Figure 4:**
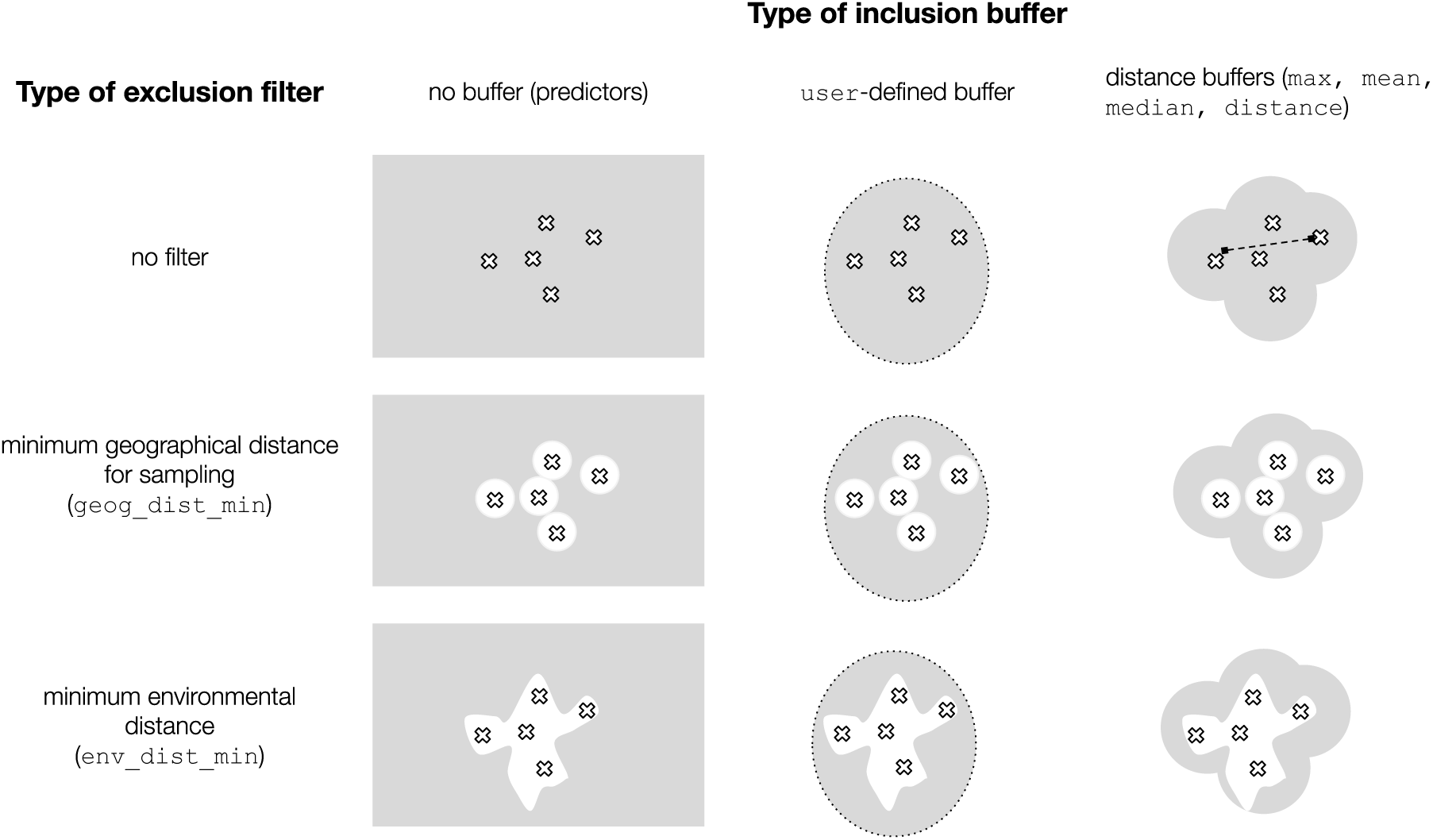
Options included in **modleR** for the sampling of pseudoabsence points

For the inclusion buffers, the user may supply a shapefile with the desired calibration area (`buffer_type = “user”`). This is the option for workflows where the design of the M portion of the BAM diagram (Barve et al., 2011; Peterson et al., 2011) is obtained externally. The inclusion buffer area can also be calculated as a distance around the occurrence points, either as an absolute numeric distance (in the units of the environmental variables, ex. `buffer_type = “distance”` and `dist_buf = 2` mean a buffer of 2 degrees around the occurrences), or using pairwise distances between points (parameter `buffer_type`, options “mean”, “maximum” and “median”, respectively).

For the exclusion filters, the minimal distance in the geographical space is controlled by parameter `min_geog_dist` and expressed in the units of the predictor variables raster. In the environmental space (env_filter = TRUE), the euclidean environmental distance to the occurrences will be calculated, either as the distance of each pixel in the predictor variables to the environmental *centroid* of the occurrence dataset (the multivariate median of the environmental values for each occurrence, `env_distance = “centroid”`), or as the minimum distance of each pixel in the predictor variables to any occurrence point (`env_distance = “minimum”`). The scale of the environmental distance is inversed to transform it into similarity, so it varies from large negative values, corresponding to high values of distance and low suitability, to 1, corresponding to zero distance and total similarity. The range of values depends on the predictor variables and their extent. Parameter `min_env_dist` will be used to set the distance that should be omitted from pseudoabsence sampling, expressed in quantiles of the overall distance range (e.g. 0.05 is 5% of the distance range -to either the centroid or to any occurrence point- and will omit the pixels within that distance).

The inclusion buffer and the exclusion filters can be applied independently or simultaneously (Figure 4), and function `setup_sdmdata()` will sample the pseudoabsences in the resulting calibration area, using `randomPoints()` function form **dismo** package.

### 3.3 Variable selection

An *a priori* selection of abiotic variables is emphatically recommended by many authors (Peterson et al., 2011; Fourcade et al., 2018), in order to grasp the relevant dimensions of the ecological niche that affect species distribution. An additional question is whether such variables are correlated, and how this could result in model overfitting (Braunisch et al., 2013). In **modleR**, the maximal correlation between the predictor variables within the calibration area can be controlled by setting `select_variables = TRUE`. Function `setup_sdmdata()` will sample a percentage of pixels in the calibration area (`sample_proportion`, from 0 to 1), calculate a correlation matrix and use `findCorrelation()` function from R package **caret** (Kuhn, 2018), to retain the highest quantity of variables that have pairwise correlations under a user-defined value (parameter cutoff). This is an automatic approach that may remove significant variables, and it does not substitute the need for a selection based on ecological meaningfulness.

### 3.4 Data partitioning into training and test sets

During data setup, occurrence and pseudoabsence data are partitioned into training and test sets, by bootstraping or crossvalidation. Parameter `partition_type` can receive either `“bootstrap”` or `“crossvalidation”` values. Crossvalidation is controlled by parameters `cv_n` for the number of runs and `cv_ partitions` for the number of partitions for each run. Bootstrap is controlled by parameters boot_n for the number of runs and boot_proportion for the proportion of presences and absences in the dataset that will be used as training data.

### 3.5 Data setup input and output

The data setup phase receives an occurrence dataset, with at least the columns for latitude and longitude values, and a RasterStack of predictor variables. The species name must be provided, and if column names are other than “lat” and “lon”, arguments lat and lon must be specified.

At the end of the data setup step of the workflow, a `sdmdata.csv` file is created with the information required for running the models. In addition to the file `sdmdata.csv`, a metadata.csv file with the metadata of the current modeling round, and a `sessionInfo.txt` file with the packages loaded and their version and origin will be created. Optionally, a .png image of the data can be created (if parameter `png_ sdmdata = TRUE`), and the final calibration area for pseudoabsences can be saved as a raster to the disk (with write_buffer = TRUE). The structure of the sdmdata data frame reflects all setup decisions as follows:

- The first `n_occs` rows (or the corrected number after data cleaning) correspond to presences and the last n_back rows (or the corrected number due to available sampling pixels) to the sampled pseudoabsences
- The first columns hold the data partition vectors. These can be named `“boot_1”` to `“boot_n”` in the case of bootstrap, `“cv_0”` in the case of one-time cross-validation or `“cv_1”` to `“cv_n”` in the case of repeated cross-validations. In the case of bootstrap, each column consists of 0 and 1 to divide into training and test set, and in the case of cross-validation, of numbers from 1 to n, where n is the number of partitions. The coding for a zero in bootstrap cases indicates internally that this column will be run only once, while in the case of cross-validation each column is run once for each group (coded as a non-null number).
- The next column is a vector of presences and absences, named `pa`, with `n_occs` presences coded as 1 and n_back pseudoabsences coded as 0. The combination of each data partition column and this presence/absence column forms the different partitions into the training and test sets.
- The next two columns have the longitude and latitude data.
- The last columns correspond to the environmental data associated to each coordinate.

As explained before, if the user cleans the biotic data, some lines (occurrences) will be removed from this basic table. Likewise, if the user asks for variable selection according to their correlation, the output sdmdata table may have less columns than the original dataset. The resulting data frame will be saved to the HD and read by the next functions, `do_any()` or `do_many()`.

### 3.6 Data setup optimization

In workflows where the same parametrization options are passed to different algorithms, function `setup_sdmdata()` will check if there is a previous folder structure and a `metadata.csv` file. If the metadata in disk match the current function call and the folder names are unchanged, the function will skip the data setup part in order to avoid repeating the data partitioning process. This optimizes parallelization and disk usage, especially during the pseudoabsence sampling phase. If a previous metadata file is found but it has different metadata (i.e. there is an inconsistency between the existing metadata and the current parameters), it will run the function with the new parameters and overwrite metadata and sdmdata files.

However, this check does not guarantee that all data in the same folder come from the same parametrization options, since different modeling rounds can create different files and *older files are not deleted*. If an experimental approach will be taken and multiple setups will be tried by the user, it is recommended that any changes in parametrization come with a change in the destination folder and a manual check of the preexisting files in disk.

## 4 Model fitting and projecting: functions do_any() and do many()

Functions `do_any()` and `do_many()` create a model per partition per algorithm (a partition), and save them into the `partitions` subfolder. While `do_any()` performs modeling for one algorithm at a time –chosen by using parameter `algorithm–, do_many()` can select multiple algorithms, with `TRUE` or `FALSE` statements. `do_many()` is just a wrapper that will call several instances of `do_any()`. The user may choose which one to choose, but for parallelization by algorithm it may be better to call `do_any()` individually.

### 4.1 Algorithm implementation

The available algorithms have been wrapped from R packages commonly used in the field of ecological niche modeling:

- bioclim, maxent, Mahalanobis distance (`mahal`), `domain`, and Boosted Regression Trees (`brt`) come from the **dismo** package (Hijmans et al., 2017). Here, since Mahalanobis distance tends to have large negative values, we set up a hard lowest limit corresponding to the Lowest Presence Training value (**dismo** threshold `no_omission`) and scaled the resulting interval from 0 to 1. BRT are fit using the default options of function `gbm.step()` (Hastie et al., 2001; Elith et al., 2009), except for `n.minobsinnode = 5`. The rest of the **dismo** algorithms are implemented using the default parameters, including `maxent`.
- The **maxnet** package implementation of maxent (`maxnet`) has been included as well, as it does not depend on package rJava (Urbanek, 2019) for running.
- Two implementations of Support Vector Machines (SVM), come from packages `kernlab` (Karatzoglou et al., 2004) and e1071 (Meyer et al., 2017). We call these `svmk` and `svme` respectively. `svmk` is fit using the default parameters for function `svmk()` in **kernlab**, and `smve` is implemented by using a `best.tune()` approach. The latter function does not always converge on the first run, so `do_any()` will automatically retry the fit up to 10 times before returning an error. In our experience this happens very infrequently and only needs a second run to converge –or it never does.
- GLM (`glm`) comes from **base** R, and was implemented using a stepwise model selection approach (function `step()`, also from **base** R) in both directions (forward and backward), considering all possible combinations of predictor variables. When projected to the geographical space, a parameter `type = “response”` is used to return values in the scale of the response variable.
- Random Forests (`rf`) come from package **randomForest** (Liaw and Wiener, 2002). It is executed by function `tuneRF()`, that finds a best value for `mtry` (the number of variables randomly sampled as candidates at each split) and returns the best model fit with this value (parameter `doBest = TRUE`).

We aim to improve the capacity of fine-tuning these algorithm implementations in the future, to facilitate tests using different sets of parameters. This is especially important for `maxent`, for which regularization parameters, response type curves and type of extrapolation (clamping or not) can affect significatively the output of the results (Elith et al., 2011; Phillips et al., 2006; Cobos et al., 2019), and for which an approach of model selection using AIC can be applied (Muscarella et al., 2014) In the current version, algorithm parametrization can be modified by the user directly inside the source code. Finally, for statistical algorithms that require presence and absence data for fitting –brt, random forests–, an additional option to equalize the number of pseudoabsences and the number of presences can be applied via `equalize = TRUE`, following Barbet-Massin et al. (2012).

### 4.2 Model evaluation

After model fitting, `do_any()` and `do _many()` execute the evaluation functions available in **dismo**, such as `evaluate()` and `threshold()`, and organize the evaluation statistics into two tables that are written into the hard disk.

The first table corresponds to the output of function `evaluate()` from package **dismo**, that calculates 1) the confusion matrices (true positives, false positives, false negatives and true negatives), 2) the relevant evaluation statistics along the whole vector of possible thresholds (for further details we refer the reader to **dismo** documentation).

Other than **dismo** performance metrics, we have included the calculation for the True Skill Statistic (Allouche et al., 2006), the F-score (also known as Sorensen’s dissimilarity score) (Powers, 2011) and Jaccard’s dissimilarity score (Leroy et al., 2018) for each threshold. The suffix of this first table when it is written to disk is `eval_mod`.

The second data frame summarizes 1) the relevant thresholds that maximize the performance metrics (see **dismo** documentation), and the corresponding maximized values of TSS, Cohen’s Kappa ((Cohen, 1960)), F-score and Jaccard 2) the performance metrics that do not depend on a threshold, such as AUC (from **dismo**) and partial ROC (Peterson et al., 2008), as implemented by package **kuenm** (Cobos et al., 2019), and their significance. This second table also indicates the performance metrics relative to the threshold specified by the user (parameter dismo_threshold in the function and column with the same name in the table).

### 4.3 Projection to other datasets

By default, `do_any()` projects the fit model unto the prediction variables (named “present” for convenience, although the predictor variables used for model fitting can correspond to any moment). It can also project the models to other predictor variable sets, representing a shift either in time or in space.

For this, the user has to setup a folder with the environmental variables, with one subfolder per projection, and make sure that the names of the variables in these folders match exactly the variables in the original environmental variables dataset. This directory should contain only raster files in their corresponding subfolder. The subfolders can hold complete datasets (ex. bio 01 to bio 19 in Worldclim), and the function will only use the rasters that were selected during data setup.

Functions `do_any()` and `do_many()` will read this folder and perform all projections when `project_model = TRUE`, and the relative or absolute location of the variable folder is passed to parameter `proj_data_folder`. For reproducibility purposes we recommend that all paths be relative.

### 4.4 do_any() and do_many() input and output

The input for `do_any()` and `do_many()` are the species name, the location of the models_dir folder previously created by `setup_sdmdata()`, and, optionally, the location of the folder with the environmental variables for projection (`proj_data_folder`). Internally, the functions will read the sdmdata.csv file created by `setup_sdmdata()`.

Regarding the functions outputs, at the end of a modeling round the partition folder will contain:

- A .tif file for each partition. If specified by the user, (parameter write_bin_cut), a binary version cut by the specified `dismo_threshold`, and a “cut” version –obtained by keeping the continuous values of the raw model above this threshold and setting all the areas below the threshold as 0– can also be created and written to disk.
- Optional .rda files with the fitted model objects for each partition and algorithm. These files can be saved by using parameter `write_rda = TRUE` and their names contain the species name, partition number and algorithm (e.g., `Abarema langsdorffii_1_1_bioclim.rda`). These .rda objects can be loaded to the workspace with function `load()`, if needed to apply other functions within R
- Figures in .png to explore the results readily, without reloading them into R or opening them in a SIG program. The creation of these figures can be controlled with parameter `png_partitions`.
- A data frame with the evaluation data for each partition for all considered thresholds, that corresponds to the output of `evaluate()` function in **dismo**, and the additional performance metrics. This is a csv file with the name containing the species name, partition number and algorithm (e.g., `eval_mod_Abarema_langsdorffii_1_1_bioclim.csv`).
- A data frame with the summary of the evaluation data for each partition, including the important thresholds (output from function `threshold()`), the threshold-independent metrics, and the specific metrics for the selected given threshold. This is a csv file with the name containing the species name, partition number and algorithm (e.g., `evaluate_Abarema_langsdorffii_1_1_bioclim.csv`).
- If any projection was executed, additional subfolders with the projection results will be created (Figure 3).
- Most importantly, `metadata.csv` and `sessionInfo.txt` files will record all the relevant parametrization options and packages that were used during the last modeling run.

The final models per algorithm will be created in the next step of the workflow.

## 5 Partition joining: final_model()

Function `final_model()` joins the partition results into one model per species per algorithm. This is one of the least documented steps in ENM workflows, that usually skip the partition joining phase and discuss algorithm comparison or consensus. The most straightforward way to build such a model is to obtain a central measure from these partitions (e.g. a mean or a median) and ideally some variation metric, and to examine the related mean performance metrics.

However, there is no consensus about the final way an ENM should be presented and depending on the application the user may want a continuous output, or a binary outcome, or understanding the degree of agreement between individual partitions. In response to this, `final_model()` has the following outcome possibilities. They all come from the raw continuous partitions.

- `raw_mean` is the mean of the raw partitions and therefore has a continuous scale
- `raw_mean_th` is raw_mean cut by the *mean threshold value that maximizes a selected performance metric* raw_ mean_th, and is a binary model
- `raw_mean_cut` recovers the original continuous values of raw_mean above this mean threshold. It has a continuous scale above said threshold and zeros below it
- `bin_ mean` is the mean of the binary partitions (created by cutting the raw partitions by their individual threshold, also set by parameter raw_mean_th). If there are n partitions, this results in a discrete categorical scale that varies from 0 to 1 in 1/n intervals. The scale reflects the number of partitions that predict the species presence on each pixel, i.e. the degree of consensus between the partitions
- `bin_consensus` cuts `bin_mean` by a specific consensus level to create a binary outcome, set by parameter `consensus_level` (e.g. 0.5 are areas predicted by half of the partitions, a majority consensus).

It is helpful to see the sequence from `raw_ mean` to `raw_mean _cut` as a “take the mean first, make binary later” logic, and the sequence from `raw_mean` to `bin_consensus` as a “make binary first, take the mean later” logic (Figure 5). These options can be passed to parameter `which_final`, either individually or as a character vector e.g. c(`“raw_mean”, “bin_consensus”`).

**Figure 5:**
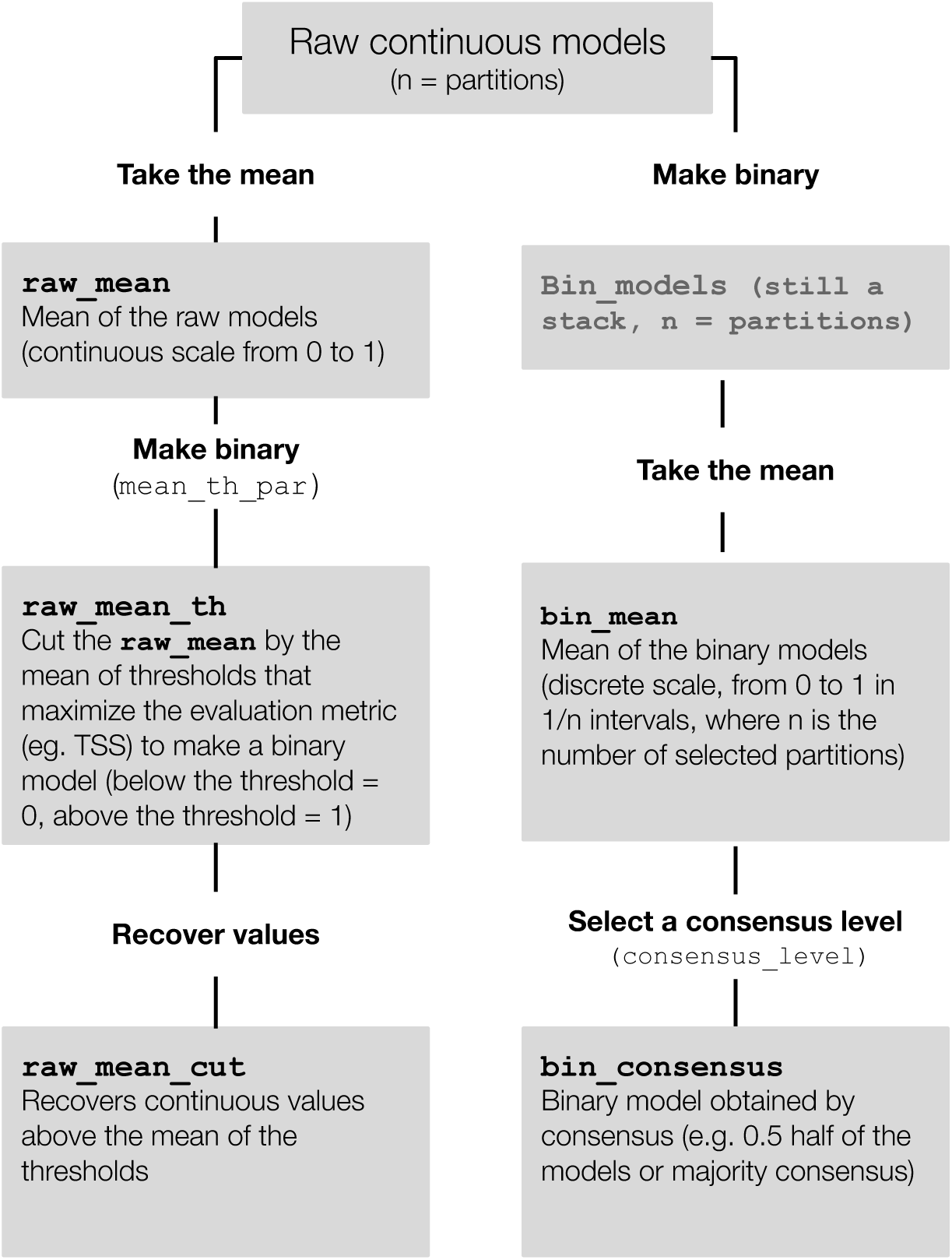
Joining partition models into final models in **modleR**

### 5.1 Uncertainty between partitions

In **modleR**, variation measures are implemented in this step and the following (see section 6). Variation between partitions should be assessed and understood before reaching to conclusions (Pearson et al., 2006; Diniz-Filho et al., 2009) in ENM, and recently many studies have focused on the development and analysis of variation metrics.

In **modleR**, the range of the adequabilities (maximum - minimum) for each pixel represents the variation between partitions. Since the number of partitions (and later, of algorithms) tends to be low and the distribution of adequability data is not expected to be normal, we opted out of calculating other metrics, like the standard deviation. The user however may extract any variation value directly from the rasterStack saved in disk from the last step. Parameter `uncertainty = TRUE` controls the creation of this uncertainty map.

### 5.2 Other parametrization options in final_model()

As explained in section 2.2, the folder where final models are saved can be renamed, by using parameter `final dir`. In a testing context, we recommend that different rounds be saved in different folders.

Parameter `algorithms` can take a character vector, e.g. c(`“bioclim”, “rf”`), to indicate specifically which algorithms should be processed. If left unspecified, models from all algorithms present in the partition folder will be used.

Parameter `proj_dir` can be passed to the function in order to perform this partition joining to projections other than `“present”`. In this case the average threshold will come from the present statistics table. The only way to execute this function to projected data while avoiding any threshold that comes from the present is `raw mean`.

### 5.3 final_model() input and output

Function final_model() reads the partitions on the hard disk, so no object in the workspace is needed to run it. At the end of the third step of the workflow, the `final_models` folder will contain:

- A .tif file for each different final models for 1) the selected algorithms and 2) the specified outputs (`which_model`)
- A .tif file for the uncertainty, if requested by the user
- Optional figures in .png to explore the results without loading them to R or a SIG program (parameter `png_final`)
- A .csv data frame with the statistics for the partitions (summarizing the results from `evaluate()` function in package **dismo**)
- A .csv with the mean thresholds and the mean performance metrics per algorithm
- A .csv file with the metadata associated with this step
- A .txt file with the session information associated with this step

## 6 Algorithmic consensus: function ensemble_model()

The fourth and final part of the **modleR** workflow joins the different models per algorithm into an ensemble, or algorithmic consensus model (Araújo and New, 2007). As with every other step in the ENM workflows, there is no agreement on which is the best way to implement such consensus models.

The discussion about consensus models is related to the fact that along the ENM/SDM workflows there are many sources of uncertainty. Source data, sampling biases and the large amount of options in algorithm choice and parametrization accumulate and create a variety of responses that make interpretation difficult. The magnitude and relative importance of these variation sources should be understood before interpreting any ENM/SDM result. Many discussions about this step include also the projection into other datasets, that adds the different GCMs and emission representative pathways as strong uncertainty sources. These comparisons have concluded that algorithm choice and parametrization is a source of uncertainty that can be larger than the choice between GCMs or RCPs [ref]. Other workflows, such as **kuenm**, analyze the relative importance of these uncertainty sources but here in **modleR** we calculate range-based variation metrics (maximum-minimum).

We have implemented several ways to calculate the ensemble models in **modleR**, partly implemented in other works (Marmion et al., 2009; Pearson et al., 2006; Zhu and Peterson, 2017):

- `“best”` chooses the best algorithm according to a desired `performance_metric` (ex. AUC, TSS)
- `“average”` is the mean of the continuous final models obtained in the previous step as `raw_mean`
- `“weighted_average”` is the weighted mean between the continuous `raw_mean` models according to a `performance_metric` (Marmion et al., 2009)
- `“median”` is the median between the continuous `raw_mean` models
- `“frequency”` is the mean of the binary models (`raw_mean_th`), which is analogous to a frequency count. Here, the user may select which threshold they want to use, independently from the decision taken in the previous workflow step, and for this they may use parameter `dismo_threshold`
- `“consensus”` extracts a binary consensus from the frequency count (e.g. `consensus_level = 0.5` means a majority consensus)
- `“pca”` extracts the first axis of a PCA between the `raw_mean` models (Pearson et al., 2006)

In addition to these ensemble options, the uncertainty metric based on ranges (maximum - minimum values) can be written using parameter `uncertainty`. It is important to note that in spite of being widely used, this last step of algorithmic consensus can be useful sometimes but it does not necessarily perform better than individual algorithms (Zhu and Peterson, 2017).

### 6.1 Input and output for ensemble_model()

Function `ensemble_model()` reads the evaluation tables and final models on disk, so that outside parametrization, no object in the workspace is actually needed to run. Optionally, if writing png files with parameter `png_ensemble` and asking for the original occurrences to be plotted, the occurrence data frame should be provided.

As in the previous steps of the workflow, the output for ensemble_model() are the .tif files, optional .png figures and the metadata.csv and sessionInfo.txt files with the parametrization options and packages.

## 7 Optimization and flexibilization decisions

During **modleR** implementation, we have included some features that allow the users to execute projects that need higher performance. At all steps, the optional outputs such as images, binary and cut models are only created in the workspace if they are going to be written to the disk, sparing processing time. All the non-optional outputs are written to the disk and read by the functions, so they do not occupy RAM memory in the workspace from one step to the other.

The fact that the basic modeling fits one model per algorithm and species or several algorithms per species allows the user to parallelize the modeling procedure by species or by algorithm. Although **modleR** does not have internal parallelism options, a parallelization cluster can be set up within R. The simplest form of a multi-species workflow would be constituted by a loop along a vector of species names and a list of coordinates. However, more detailed parametrization may be desirable, and in this case several lists of parameters can be prepared and fed into function `clusterMap()` from package **parallel** (R Core Team, 2018), for instance. **modleR** constitutes a good testing platform, since at all steps one or some of the parameters can be modified and this does not depend on running the overall workflow repeatedly.

## 8 Next steps and further implementations

Due to the very nature of the workflow and the ecological niche modeling discipline, **modleR** is work in progress, and will be maintained and expanded depending on new theoretical developments. However, a shortlist of next steps includes several improvements such as:

- Independent in-depth parametrization for each algorithm
- Internal parallelization to optimize performance
- adapting the previous Shiny application to this new implementation of the package

## 9 Installation

Currently, **modleR** is available from GitHub ^3^ and can be installed by using the following commands:

~~~
# Without vignette
remotes::install_github(“Model-R/modleR”, build = TRUE)
# With vignette
remotes::install_github(“Model-R/modleR”,
                        build = TRUE,
                        dependencies = TRUE,
                        build_opts = c(“--no-resave-data”, “--no-manual”),
                        build_vignettes = TRUE)
~~~

The package has a vignette and a pkgdown Wickham and Hesselberth (2019) site ^4^ with the basic workflow usage information.

## 10 Conclusions

We introduced and detailed **modleR**, a four step R workflow for ecological niche modeling based on **dismo** (Hijmans et al., 2017) and other packages in the R statistical environment, directed at solving the trade-off between the common tasks that are only superficially repetitive, and the need for flexibility in decision making. The structure of this workflow includes the basic steps that an ENM project should perform and documents them thoroughly, not only by using a script-based approach, but by recording explicitly the metadata associated with each modeling step.

The parametrization options and the folder structure flexibility allow for testing different modeling options conveniently, without the need for repeating previous steps. Furthermore, the simple objects that are used as input and outputs allow for comunication with other ENM/SDM-related R packages.

We expect that this workflow can be used as a testing platform and expand with newer tools and consensus in the field and welcome users and collaborators to contribute to its development.

## 11 Acknowledgments

We thank A. Townsend Peterson for his comments and suggestions about the workflow structure, and all the users that have tested the package and helped us improve it. The authors have declared no competing interests.

## 12 Author contribution

AST, SRM, DSBR, FSMB and GG wrote the R package. AST, SRM, and MFS wrote the manuscript. MFS supervised the project. All authors reviewed the manuscript and gave final approval for publication.

## Footnotes

1 https://diva-gis.org

2 https://openrefine.org/

3 https://github.com/Model-R/modleR

4 https://model-r.github.io/modleR/index.html

## Notes

https://model-r.github.io/modleR/index.html

https://github.com/Model-R/modleR

